# Systemic exposure to COVID-19 virus-like particles modulates firing patterns of cortical neurons in the living mouse brain

**DOI:** 10.1101/2024.11.26.625543

**Authors:** Aniruddha Das, Jacob Icardi, Julie Borovicka, Sarah Holden, Henry F. Harrison, Alec J. Hirsch, Jacob Raber, Hod Dana

**Author notes:** equal contributors (Aniruddha Das, Jacob Icardi).

## Abstract

Severe Acute Respiratory Syndrome Corona Virus 2 (SARS-CoV-2) causes a systemic infection that affects the central nervous system. We used virus-like particles (VLPs) to explore how exposure to the SARS-CoV-2 proteins affects brain activity patterns in wild-type (WT) mice and in mice that express the wild-type human tau protein (htau mice). VLP exposure elicited dose-dependent changes in corticosterone and distinct chemokine levels. Longitudinal two-photon microscopy recordings of primary somatosensory and motor cortex neurons that express the jGCaMP7s calcium sensor tracked modifications of neuronal activity patterns following exposure to VLPs. There was a substantial short-term increase in stimulus-evoked activity metrics in both WT and htau VLP-injected mice, while htau mice showed also increased spontaneous activity metrics and increase activity in the vehicle-injected group. Over the following weeks, activity metrics in WT mice subsided, but remained above baseline levels. For htau mice, activity metrics either remain elevated or decreased to lower levels than baseline. Overall, our data suggest that exposure to the SARS-CoV-2 VLPs leads to strong short-term disruption of cortical activity patterns in mice with long-term residual effects. The htau mice, which have a more vulnerable genetic background, exhibited more severe pathobiology that may lead to more adverse outcomes.

## Introduction

Infection by Severe Acute Respiratory Syndrome Corona Virus 2 (SARS-CoV-2) or one of its newer derivatives may cause systemic harm to the human body, including the brain. Reported brain-related pathologies include changes to synaptic homeostasis^1^, fusion of neurons to neurons and/or glial cells that compromise neuronal firing^2^, as well as reduced burst activity of neurons^3^. Such modified neuronal physiology may lead to behavioral alterations and cognitive deficits, which may be of major concern due to the wide spread of the virus. Moreover, at-risk populations due to age, health condition, and/or genetic risk factors of age-related cognitive decline and neurodegenerative disorders, such as in the elderly or patients with Alzheimer’s disease (AD), may suffer more from COVID-19 complications or long-term effects than the general population^4-6^.

The infectious nature of SARS-CoV-2 makes comprehensive studies with it challenging, which has led to a search for appropriate models that would allow conducting experiments in a safer environment. Such an alternative model is using virus-like particles (VLPs), which express the SARS-CoV-2 essential proteins without the replicating viral RNA^7-9^. The SARS-CoV-2 VLPs allow studying their effects on the host organism without the associated risk of being exposed to the virus itself. Previous works with VLPs showed their usefulness for testing how different SARS-CoV-2 mutations like Delta and Omicron affected the virus infectivity in the presence of vaccines^10^, or to study the role of different protein mutations on virus infectivity^11^.

Exposure to the SARS-CoV-2 virus, or to its VLPs, is expected to cause a systemic immune response with multiple potential outcomes, including activation of glial cells in the central nervous system and initiation of inflammatory responses. Therefore, such exposure would also affect neurons in the brain and may modulate their activity patterns^12-14^. The immune response is expected to decrease over time, but it remains unclear whether there would be long-term changes in brain functionality. Therefore, proper assessment of the brain may benefit from the use of longitudinal methods, which will allow for studying the progression of pathology over time.

Here, we conducted repeated recordings of neurons in the mouse motor and somatosensory cortices using two-photon laser scanning microscopy (TPLSM)^15^. Mice were first injected with VLPs to assess the dose-dependent response of their immune systems. Then, we recorded spontaneous and stimulated activity patterns from cortical neurons before and after exposure to a selected dose of VLPs. We also compared the effects of VLP exposure on mice that express the wild-type human tau protein (htau mice), which is associated with AD-related pathology^16,17^. Our findings show strong short-term effects of exposure to VLPs on the brain activity metrics, with residual long-term effects and increased vulnerability of the htau mice.

## Results

### Finding the optimal VLP dose to achieve a robust immune response

VLPs were generated, similar to our previous report^9^, to express the SARS-CoV-2 nucleocapsid (N), membrane (M), and envelope (E) structural proteins together with the spike protein (S), allowing us to perform SARS-CoV-2-related studies without replicating virus and outside of a BSL-3 facility. Mice were injected (*n =* 19 mice; intraperitoneal (IP)) with different doses of VLPs or vehicle solutions. One hour later, blood samples were collected and the levels of Chemokine (C-C motif) ligand 7 (CCL7) and corticosterone hormone in the plasma were analyzed to assess activation of the immune system by exposure to the VLPs. VLP injection showed a significant dose-dependent effect on plasma levels of CCL7, with the lowest levels being from mice treated with 1 μg VLPs versus vehicle (Fig. 1A). In addition, the injection of 1 μg VLPs significantly increased corticosterone levels 1 hour after VLP injection, while 0.3 μg VLPs showed no significant change compared to vehicle injection (Fig. 1B).

**Figure 1.**
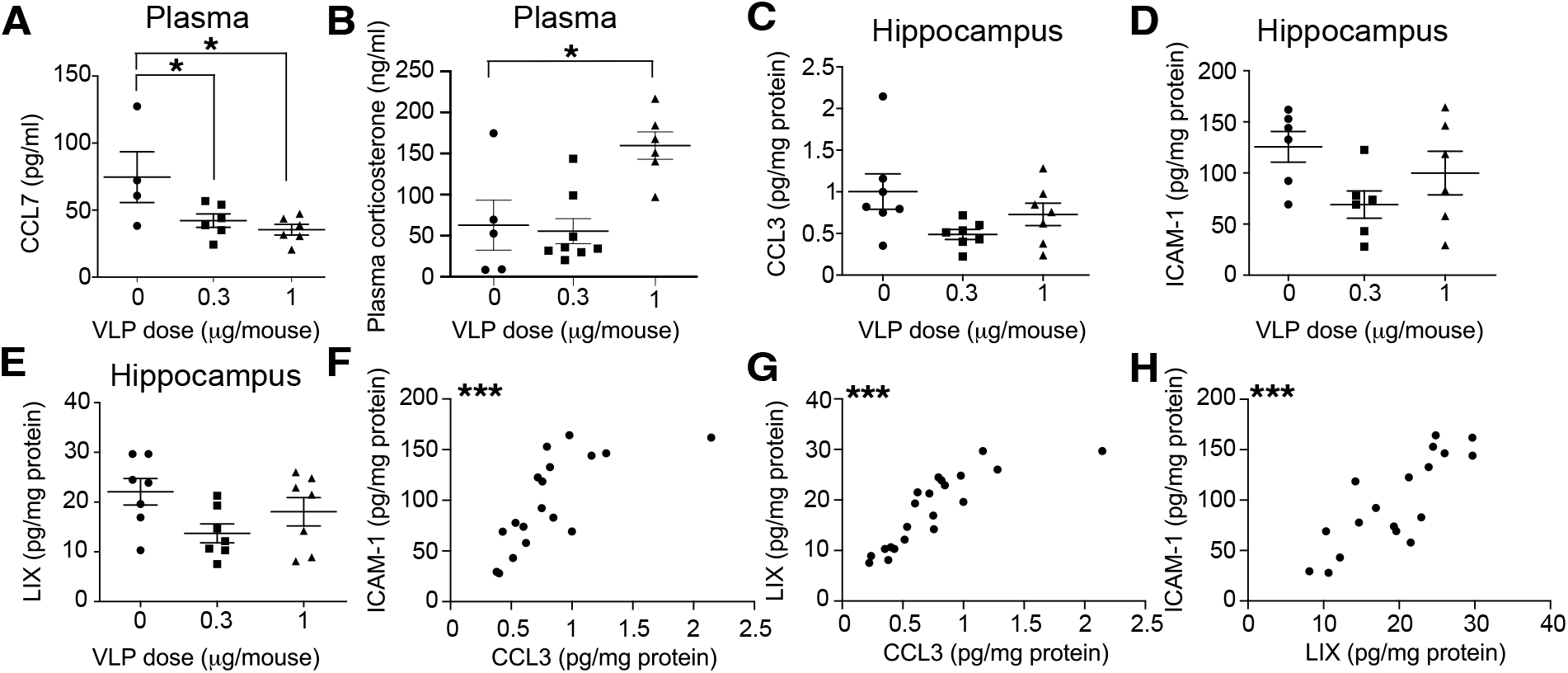
Selection of VLP dose to affect the mouse immune system. **A**. VLP injections showed a significant dose-dependent effect on the plasma levels of CCL7 (*F* = 4.703, p = 0.0291, all tests in A-E are one-way ANOVAs. All *post hoc* comparisons were conducted using Dunnett’s correction for multiple comparisons). **B**. Injection of 1μg VLPs caused a significant increase in plasma corticosterone levels, while injection of 0.3μg caused no significant change compared to the vehicle injection (*F* = 8.512, *p* = 0.003). **C**. There was a trend towards a dose-dependent effect of VLPs on hippocampal CCL3 levels (*F* = 2.976, *p* = 0.07). **D**. There was a trend towards a dose-dependent effect of VLPs on hippocampal ICAM-1 levels (*F* = 2.798, *p* = 0.09). **E**. There was a trend towards a dose-dependent effect of VLPs on hippocampal LIX levels (*F* = 2.807, *p* = 0.09. **F**. Positive correlation between hippocampal levels of ICAM-1 and CCL-3 (*r* = 0.7115, *p* = 0.0009, Pearson’s correlation). **G**. Positive correlation between hippocampal levels of LIX and CCL-3 (*r* = 0.8324, *p* < 0.0001, Pearson). **H**. Positive correlation between hippocampal levels of ICAM-1 and LIX (*r* = 0.8184, *p* < 0.0001, Pearson). *n* = 19 mice; *, *p* < 0.05; **, *p* < 0.01; ***, *p* < 0.001.

Twenty-four hours after the VLP injection, all mice were euthanized, and hippocampal tissue was extracted and processed. The concentrations of three chemokines that indicate activation of the immune system showed dose-dependent trend toward significance, with positive correlations between the measured proteins. The concentration of Chemokine (C-C motif) ligand 3 (CCL3) was decreased following injection of 0.3 and 1 μg VLPs. Similarly, we found a decrease in the concentration of Intercellular Adhesion Molecule 1 (ICAM-1, also known as Cluster of Differentiation 54, CD54), which plays an important role in mediating immune and inflammatory responses and is known for regulating leukocyte recruitment from circulation to sites of inflammation^18^. Finally, the concentration of the murine LPS-induced CXC chemokine (LIX), which is a neutrophil-chemoattractant CXC chemokine and is involved in recruitment of neutrophils to sites of inflammation^19^, was also decreased following injection of 0.3 and 1 μg of VLPs (Fig. 1C-E). Notably, there was a positive correlation between hippocampal levels of ICAM-1 and CCL3, LIX and CCL3, and ICAM-1 and LIX (Fig. 1F-H; *r* = 0.71, 0.83, and 0.82, respectively; Pearson’s correlation). Based upon these findings, the 0.3 μg dose was selected for the cortical activity recording experiments described below.

### Monitoring baseline cortical activity reveals differences between WT and htau mice

Spontaneous and stimulated activity of Layer II/III neurons in the primary somatosensory and motor cortices (S1 and M1, respectively) were recorded for both WT and htau mice (Fig. 2A-C). The two genotypes showed different brain activity patterns. The fraction of neurons that fired spontaneously and their median firing rates were significantly higher in WT mice than in htau mice (Fig. 2D). In contrast, the fractions of neurons that significantly increased their firing during paw stimulation or that maintained significantly higher jGCaMP7s fluorescence levels after the end of the stimulation period than their baseline fluorescence levels (tuned or sustained activity neurons, respectively), as well as the fraction of neurons that exhibited both tuned and sustained activity were significantly higher in htau than WT mice (Fig. 2E). Finally, in WT mice, but not htau mice, the median spontaneous firing rate of S1 neurons was significantly higher than that of M1 neurons (Fig. 2D), which was also the case for the fraction of S1 neurons that were tuned to paw stimulation or showed both tuned and sustained activity (Fig. 2E). Due to these fundamental differences in brain activity patterns among the two mouse lines, the analysis of VLP exposure effect on each line was performed separately in the following sections.

**Figure 2.**
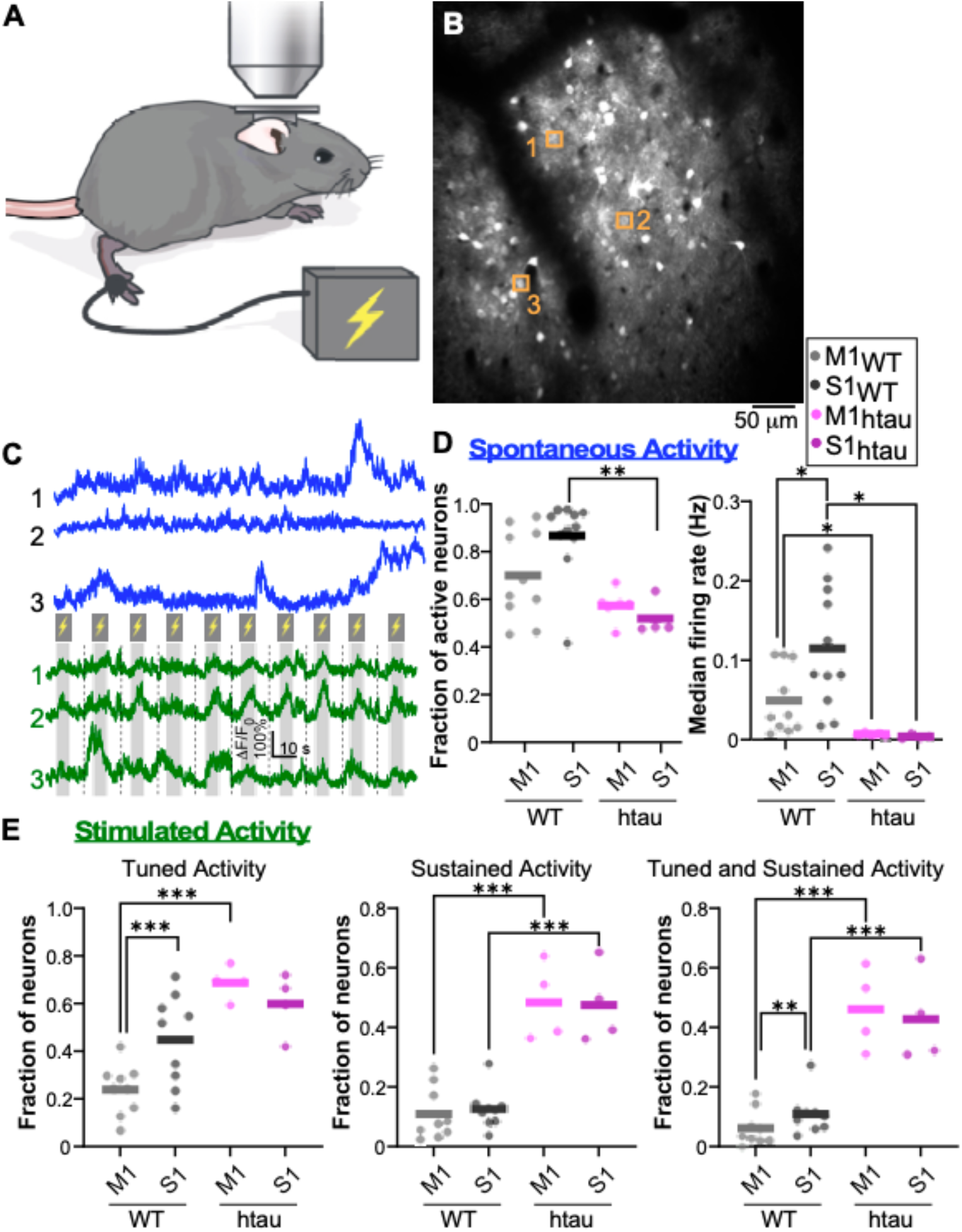
Recording of baseline spontaneous and stimulated activity in WT and htau mice. **A**. Schematic illustration of the experimental setup. Mice were lightly anesthetized, sedated, and received trains of electric stimuli to their hindlimbs while neuronal activity from their motor and somatosensory cortices was recorded using two-photon microscopy. **B**. Example image of S1 neurons expressing jGCaMP7s. **C**. Examples of spontaneous and stimulated activity traces (blue and green traces, respectively) from the cells indicated by orange boxes in panel B. Gray bars and lightning signs indicate the times where the paw stimulation was given to the mouse, while dashed lines indicate the separation between stimulation cycles. **D**. Comparison of the fraction of neurons that fired action potentials during the spontaneous activity recordings in S1 and M1 out of all recorded neurons in the same brain region (left, active neurons), and the median firing rate of the active neurons (right). Each dot shows data from one mouse, horizontal bars are the mean per brain region. The fraction of active neurons was significantly higher in the S1 region of WT mice than that of htau (*p* = 0.0014, unpaired Student’s t-test; 0.87±0.16 for WT S1, 0.7±0.19 for WT M1, 0.52±0.08 for htau S1, and 0.58±0.08 for htau M1, mean ±std). The right panel shows the median firing rate, where each dot is the median across all the recorded active neurons from the same brain region of one mouse, while horizontal bars are the regional means. WT median firing rates were significantly higher than those in htau for both S1 and M1 regions (*p* = 0.015 and 0.0486, unpaired Student’s t-test, respectively; 0.12±0.08 for WT S1, 0.049±0.04 for WT M1, 0.004±0.003 for htau S1, and 0.007±0.002 for htau M1, mean±std). In addition, the median firing rate in S1 of WT mice was significantly higher than that in M1 (*p* = 0.015, paired Student’s t-test). **E**. Comparison of the stimulated activity metrics shows a significant increase in htau vs. WT fraction of M1 neurons that increased their activity during paw stimulation (tuned activity), after the stimulation (sustained activity), or showed both tuned and sustained activity (*p* < 0.001 for all comparisons, unpaired Student’s t-tests; 0.69±0.07 vs. 0.24±0.11 for tuned activity, 0.48±0.13 vs. 0.11±0.09 for sustained activity, and 0.46±0.14 vs. 0.06±0.06 for both). A similar comparison for the S1 region showed that htau mice had a higher fraction of sustained activity as well as both tuned and sustained activity neurons (*p* < 0.001 for both comparisons, unpaired Student’s t-tests; 0.6±0.13 vs. 0.45±0.19 for tuned activity, 0.48±0.13 vs. 0.13±0.07 for sustained activity, and 0.43±0.15 vs. 0.11±0.07 for both). In addition, for WT mice, a higher fraction of S1 than M1 neurons was tuned and showed both tuned and sustained activity (*p* = 0.0003 and 0.0093, paired Student’s t-test). Data from *n* = 11 WT and 5 htau mice are include in D and from *n* = 9 WT and 4 htau mice in E; 49-708 and 59-1172 neurons were recorded to calculate each data point in D and E, with median values of 355 and 512 neurons, respectively; *, *p* < 0.05; **, *p* < 0.01; ***, *p* < 0.001.

### Short-term increases in firing properties following exposure to SARS-CoV-2 VLPs

Following the recordings of baseline neuronal activity, both WT and htau mice were randomly assigned to 2 groups and received a single IP injection of 0.3 μg VLPs or the same volume of vehicle solution (VLP group: 6 WT mice, 3 males and 3 females, and 3 htau mice: 2 males and 1 female; vehicle group: 6 WT mice, 3 males and 3 females, and 3 htau mice: 2 males and 1 female). Recording of brain activity started one hour after the IP injection and continued for an additional six weekly recordings. When analyzing the changes in brain activity during baseline recording, the day of injection and the following week (short-term effects) in WT mice, there was a respective mean increase of 420% and 145% in M1 and S1 stimulus-evoked response magnitudes following VLP exposure, which was different across the VLP and vehicle groups, indicating a short-term change in neuronal firing properties (Fig. 3A). Comparing the short-term changes in the fraction of tuned neurons in each brain region showed differences across the vehicle and VLP groups and an increase of 30-80% over time for the VLP group (Fig. 3B). Similarly, there was a significant short-term increase of 90-125% in the fraction of neurons with sustained activity and an increase of 95-240% in the fraction of neurons with both tuned and sustained activity for the VLP group compared to baseline. The magnitudes of these increases were generally higher in the M1 region than the S1 region, which started with higher baseline levels (Fig. 3B-D).

**Figure 3.**
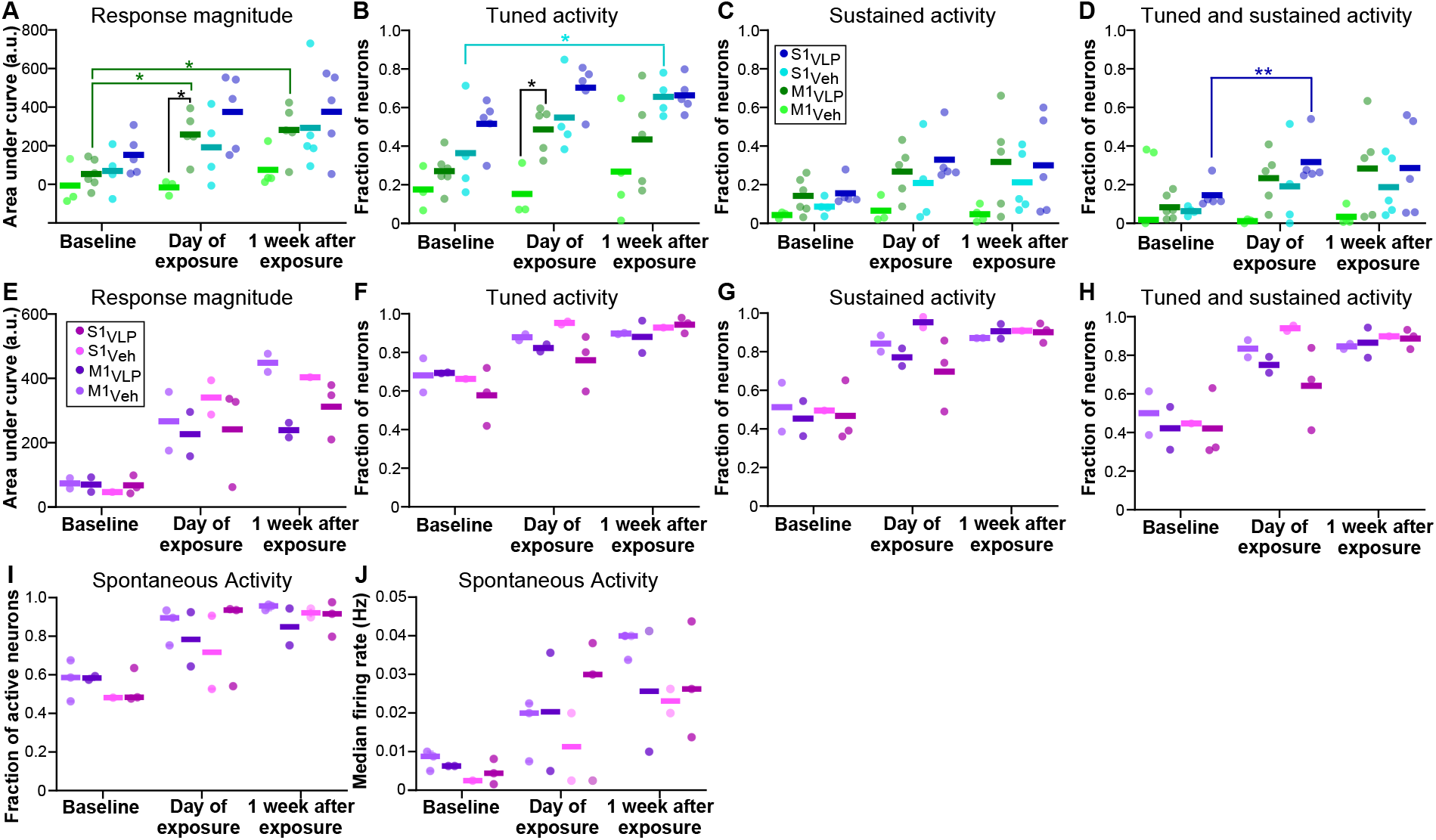
Short-term changes in neuronal firing properties following exposure to VLPs. **A**. Repeated recordings from the same mice before, on the day of VLP/vehicle injection, and in the following week, showed a significant short-term increase in the area under the curve of the measured fluorescence response magnitude to hindlimb stimulation on the exposure day (baseline levels: 53.98±72.01 in M1 and 152.69±105.32 in S1; exposure day levels: 259.1±118.81 in M1 and 375.12±193.94 in S1; arbitrary units, mean±std), a significant difference between the VLP and vehicle groups, as well as an increase the response magnitude a week after VLP exposure (group effect: F(3,16)=3.989, *p* = 0.0268; time effect: F(1.457,18.94)=12.72, *p* = 0.0008; all tests in A-J used mixed-effect models with restricted maximum likelihood. Tukey Honest Significant Difference *post hoc* comparisons were used in A-D; Data from *n* = 4-6 mice were used for each group A-D: 6 mice were included in the M1_VLP_ group, 5 in S1_VLP_, 4 in M1_Veh_, and 5 in S1_Veh_; 23-708 neurons were recorded to calculate each data point in A-D, with a median value of 188 neurons. **B**. We found a significant increase in the fraction of tuned neurons over the time after VLP exposure and a significant difference between the vehicle and VLP groups (group effect: F(3,15)=11.76, *p* = 0.0003; time effect: F(2,26)=6.079, *p* = 0.0068; M1: mean increase of 80% from 0.27±0.1 during baseline recording of the VLP group to 0.49±0.12 on the VLP exposure day, S1: 36% increase from 0.52±0.13 to 0.7±0.12 on the same days; mean±std). **C**. Same comparison of the fraction of neurons with sustained activity showed significant differences among the groups and a time effect that trended towards significance (group effect: F(3,16)=3.68, *p* =0.0345; time effect: F(1.735,22.56)=3.494, *p* = 0.0534; 124% increase in M1 from 0.14±0.09 during baseline to 0.32±0.25 one week after VLP exposure vs. 112% increase in S1, from 0.16±0.07 during baseline to 0.33±0.14 on the VLP exposure day). **D**. Same comparison of the fraction of neurons that showed both tuned and sustained activity showed significant differences between the VLP and vehicle groups, as well as significant changes over time (group effect: F(3,16)=3.66, *p* = 0.035; time effect: F(1.737,22.58)=4.467, *p* = 0.0274; 240% increase in M1 from 0.08±0.06 during baseline to 0.28±0.25 one week after VLP exposure vs. 118% increase in S1, from 0.15±0.07 during baseline to 0.32±0.13 on the VLP exposure day). For the VLP group, we noted that following VLP exposure, M1 levels increased more than S1 levels to the point that they reached similar levels, closing most of the gap that existed during baseline recording. **E**. Measuring the area under the curve of the acquired fluorescence response magnitude to hindlimb repeated stimulation of htau mice showed significant increases of 235-760% over time, but without differences across the VLP and vehicle groups (time effect: F(1.125, 4.499)=43.13, *p* = 0.0017; group effect: F(3, 5)= 0.5255, *p* = 0.6837). Data from *n* = 2- 3 mice were used each for each group in E-J: 2 mice in the M1_VLP_ group, 3 in S1_VLP_, 2 in M1_Veh_, and 2 in S1_Veh_; *Post hoc* comparisons were not conducted in E-J due to the low number of mice; 102-639 neurons were recorded to calculate each data point in E-H, with median value of 206 neurons. **F**. We found a significant increase in the fraction of tuned neurons over time without a significant difference among the groups (time effect: F(1.711, 6.845) = 16.32, *p* = 0.003; group effect: F(3, 5)=0.6852, *p* = 0.5985; 0.64±0.11 during baseline, 0.84±0.11 on VLP/vehicle exposure day, and 0.91±0.11 on the following week, mean±std, data from all recorded mice). **G**. The fraction of neurons that showed sustained activity also exhibited a significant increase over time without a significant change among the different groups (time effect: F(1.938, 7.750) = 24.86, *p* = 0.0004; group effect: F(3, 5)=0.5852, *p* = 0.6503; 0.48±0.12 during baseline, 0.80±0.14 on VLP/vehicle exposure day, and 0.90±0.04 on the following week, mean±std, data from all recorded mice). **H**. Similar to F-G, the fraction of neurons with tuned and sustained activity showed a significant increase over time without a significant difference among groups (time effect: F(1.941, 7.764)= 22.11, *p* = 0.0006; group effect: F(3, 5)= 0.6567, *p* = 0.6128; 0.44±0.13 during baseline, 0.78±0.16 on VLP/vehicle exposure day, and 0.87±0.05 on the following week, mean±std, data from all recorded mice). **I-J**. Repeated measures of changes in the spontaneous activity of neurons showed a significant increase of 64% over time in the fraction of neurons that fired action potential during the recording and of 403% in their median firing rates (from fraction of 0.55±0.08 and median firing rate of 0.006±0.003Hz during baseline to 0.91±0.07 and 0.03±0.01Hz on the week after VLP/vehicle exposure, mean±std, data from all recorded mice). There were no significant changes among the VLP and vehicle groups (time effect: F(1.448, 7.964)= 23.85, *p* = 0.0007; group effect: F(3, 6)= 0.4251, *p* = 0.7423 for I. time effect: F(1.627, 8.950)=15.05, *p* = 0.0019; group effect: F(3, 6)= 0.4024, *p* = 0.7568 for J). Each dot shows the fraction of neurons with at least one detected action potential from all recorded neurons in each brain region (I), or the median of the average firing rate across all active neurons within each region (J). 115-990 neurons were recorded to calculate each data point in I-J, with a median value of 286 neurons. Horizontal bars are the average (A-H) or median (I-J) across all data points. *, p<0.05; **, p<0.01

Measuring the same functional metrics in htau mice revealed different results. Similar to the WT mice, the neuronal response magnitude of htau mice showed a significant increase of 359% and 363% in M1 and S1 regions to paw stimulation (Fig. 3E). Following exposure to VLP, a significant increase was also detected in the fraction of tuned, sustained, and tuned and sustained neurons, which brought their values close to the maximal possible value of 1 (Fig. 3F-H. Tuned neurons: a mean increase from 0.62±0.12 during baseline recordings to 0.92±0.07 one week after VLP exposure; sustained activity neurons: from 0.46±0.13 to 0.90±0.05; neurons with both tuned and sustained activity: from 0.42±0.15 to 0.88±0.07; average on both M1 and S1 regions). Interestingly, htau mice also showed additional changes in spontaneous neuronal activity patterns that were not apparent in WT mice. There was a significant increases of 59% in the fraction of active neurons and of 412% in their median firing rates between baseline recordings and the week after VLP exposure (Fig. 3I-J; mean across M1 and S1 regions). These increases were not limited to the VLP group, and similar increases were evident in the vehicle group. No significant changes in spontaneous activity metrics of WT mice were found. These measurements indicate that systemic VLP exposure leads to short-term increases in neuronal firing properties in response to paw stimulation in both WT and htau mice; however, htau mice showed higher susceptibility to this exposure than WT mice, which was evident by the magnitude of increases in stimulated activity metrics, increases in spontaneous firing metrics, and detected increases in the vehicle group.

### Long-term changes in neuronal activity following VLP exposure

To identify and compare long-term activity changes between WT and htau mice, the 6-week post-exposure recording data were grouped into early-term (exposure day and the following week), mid-term (weeks 2-3 after exposure), and late-term measurements (weeks 4-6 after exposure; same mice and groups as for the short-term recordings). The response magnitude of WT mice significantly increased in the early time after VLP exposure and remained elevated above baseline values for the VLP group throughout the entire recording time, while the vehicle group values remained close to the baseline levels (Fig. 4A). Similarly, the fraction of tuned neurons, the fraction of sustained activity neurons, and neurons with both tuned and sustained activity of the VLP group showed an early-term increase, followed by a partial decay to an intermediate level that was higher than baseline, and with significant differences between the VLP and vehicle groups (Fig. 4B-D). Overall, WT mice showed substantial short-term increases, which reached their peaks approximately one week after the exposure to VLPs (Fig. 3A-D). Longer-term effects of VLP exposure on the brain activity were evident during the 6 post-exposure weeks that were monitored, but their magnitude were lower than the short-term effects.

**Figure 4.**
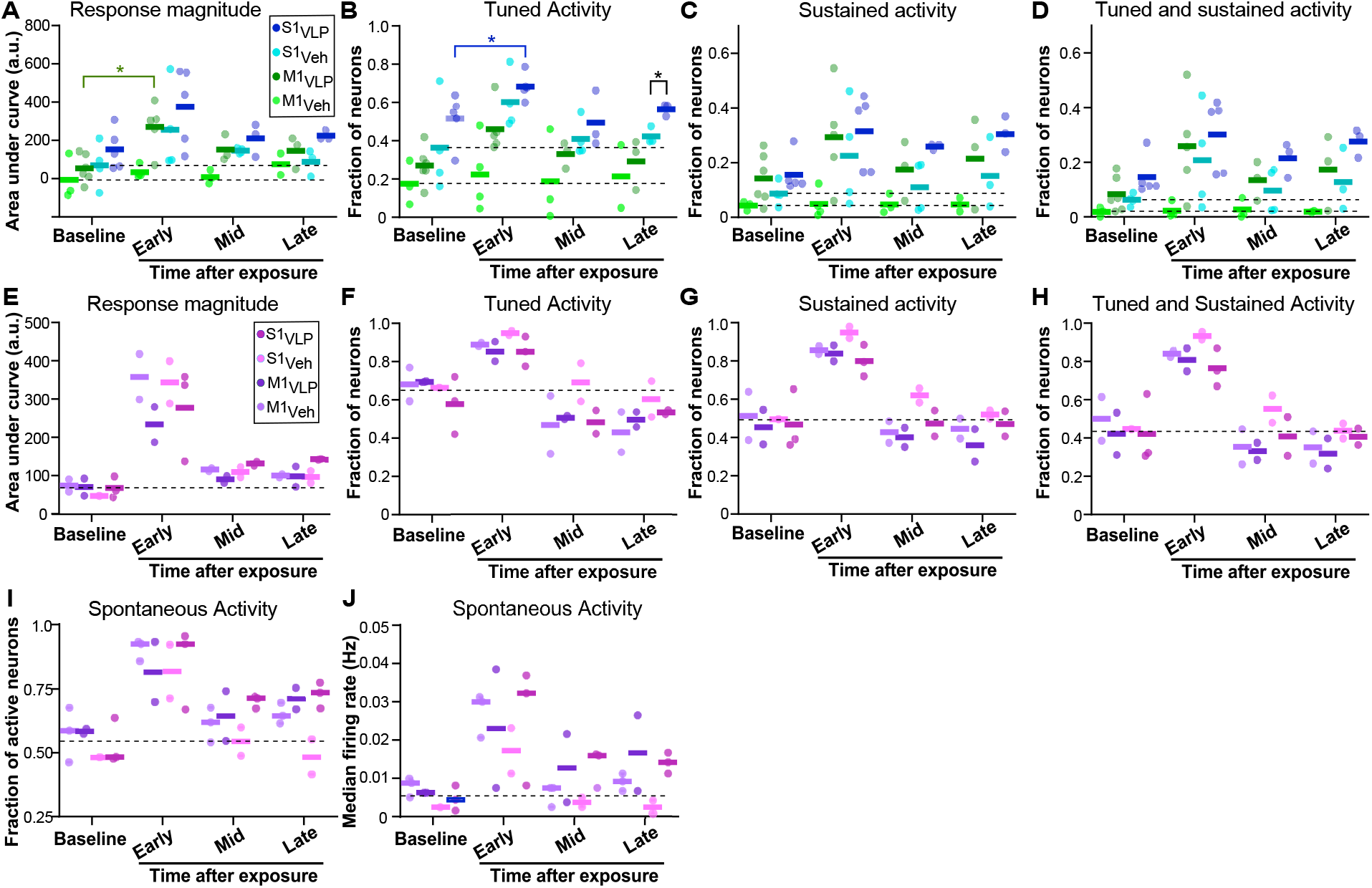
Long-term changes in firing properties following exposure to VLPs. **A**. Data were divided into baseline (pre-exposure) and post-exposure times: early (mean of same-day and the following week), mid (mean of 2-3 weeks post-exposure), and late (mean of 4-6 weeks post-exposure). The response amplitude of neurons following paw stimulation showed significant changes among the VLP and vehicle groups and over time. Interestingly, for the VLP group, there was a short-term increase following the exposure to VLP and then a decay, but the values remained elevated compared to the baseline levels (baseline levels for M1 and S1 are marked by dashed lines; group effect: F(3,15)=4.121, *p* = 0.0256; time effect: F(1.506,14.56)=8.651, *p* = 0.0055; all tests in A-J used mixed-effect models with restricted maximum likelihood. Tukey Honest Significant Difference *post hoc* comparisons were used in A-D; Data from *n* = 4-6 mice were used for each group in A-D: 6 mice in the M1_VLP_ group, 5 in S1_VLP_, 4 in M1_Veh_, and 5 in S1_Veh_; 21-708 neurons were recorded to calculate each data point in A-D, with a median value of 164 neurons. **B**. We found significant differences in the fraction of tuned neurons between the groups and over time, where after an increase in the early time after exposure, the fraction of tuned neurons of the VLP group returned to levels higher than the baseline levels (group effect: F(3,15)=10.08, *p* = 0.0007; time effect: F(1.21,11.69)=7.233, *p* = 0.0165). **C**. The fraction of neurons with sustained activity showed significant differences between groups and over time. The fraction of neurons with sustained activity in the VLP group did not return to baseline levels during the 6 weeks of recording (group effect: F(3,15)=5.371, *p* = 0.0103; time effect: F(1.767,17.08)=4.438, *p* = 0.0318). **D**. The fraction of neurons that were both tuned and with sustained firing behaved similarly to the neurons with sustained activity, where there were significant changes between the VLP and vehicle groups and over time, and the measured levels post-exposure decayed from the early-phase increase, but did not return to baseline levels (group effect: F(3,15)=5.221, *p* = 0.0115; time effect: F(1.978,19.12)=6.553, *p* = 0.0069). **E**. The response magnitude following paw stimulation in htau mice showed a large early-phase increase and then decayed to values that remained higher than baseline levels (time effect: F(1.108, 4.432) = 55.67, *p* = 0.0011; group effect: F(3, 5)=0.2317, *p* = 0.8708; Data from *n* = 2-3 mice were used for each group in E-J: 2 mice in the M1_VLP_ group, 3 in S1_VLP_, 2 in M1_Veh_, and 2 in S1_Veh_; *Post hoc* comparisons were not conducted in E-J due to the low number of mice; 67-639 neurons were recorded to calculate each data point in E-H, with median value of 190 neurons. Dashed lines in E-J show the mean baseline level of all groups. **F**. The fraction of tuned neurons showed an increase in the early phase after exposure to both VLP and vehicle and then decreased to levels lower than the baseline without a significant difference between the groups (time effect: F(1.973, 7.891) = 34.65, *p* = 0.0001; group effect: F(3, 5)=0.8176, *p* = 0.5371). **G-H**. The fraction of neurons with sustained activity and the fraction of neurons with both sustained and tuned activity showed significant temporal changes with an early-phase increase and then a return to levels close to baseline (time effect: F(1.398, 5.591) = 64.17, *p* = 0.0002; group effect: F(3, 5)=0.5852, *p* = 0.4609 for G; time effect: F(1.643, 6.574) = 60.87, p<0.0001; group effect: F(3, 5)=0.6567, *p* = 0.6241 for H). **I**. The fraction of neurons that fired at least one action potential during spontaneous activity recording showed a significant change over time, but not among groups. Interestingly, following the early-phase increase, the fraction of active neurons remained elevated for the VLP group, but returned close to baseline level for the vehicle group (time effect: F(1.51, 8.558) = 36.61, *p* = 0.0001; group effect: F(3, 6)=1.468, *p* = 0.3147). **J**. The median firing rate of active neurons during recording of spontaneous neuronal activity showed a similar pattern to I (time effect: F(1.294, 7.335) = 16.73, *p* = 0.0031; group effect: F(3, 6)=0.7859, *p* = 0.5441). 56-990 neurons were recorded to calculate each data point in I-J, with a median value of 206 neurons. *, p<0.05

For htau mice, we identified a different pattern of long-term effects following VLP exposure than in WT mice. For all measured metrics, there were no significant changes among the VLP and vehicle groups, but there were significant changes over time. The response magnitude of the recorded neurons was increased early after the exposure for both VLP and vehicle groups, and then decreased to levels that were higher than the baseline (Fig. 4E). The fraction of tuned neurons in both the VLP and vehicle groups increased early after VLP and vehicle exposure, and then decreased below baseline levels in the following weeks after exposure (Fig. 4F). The fractions of neurons with sustained activity and of neurons with tuned and sustained activity increased for both VLP and vehicle groups and then returned close to the baseline levels (Fig. 4G-H). When analyzing the spontaneous activity characteristics of htau mice, there was an increase for both VLP and vehicle groups early after the exposure. However, while there was a return of the vehicle groups to levels close to baseline, the VLP groups decreased from the early peak but remained elevated compared to the baseline levels (Fig. 4I-J).

## Discussion

This work investigates the effects of VLP exposure on neuronal activity patterns in mouse motor and somatosensory cortices in a longitudinal manner. The use of VLPs enabled studying such effects without most of the risk associated with the replicating SARS-CoV-2 virus and its derivatives, and therefore also allowed conducting more sophisticated studies, such as two-photon microscopy recording of single-neuron activity *in vivo*, that otherwise would have been more challenging to perform. This work demonstrates recording of cortical activity through an implanted cranial window for six consecutive weeks after exposure to VLPs or vehicle and identifies short- and long-term changes in various functional metrics of neurons that relate to stimulus-evoked, as well as spontaneous, activity patterns.

We tested the effects of different VLP doses on the mouse immune system and inflammatory responses, and selected the 0.3 μg/mouse dose, which elicited a strong immune response without alterations in plasma corticosterone levels. Next, we studied the effect of VLP exposure on mice and how it relates to the presence of the wild-type human tau protein. Tau is fundamentally involved in tauopathy disorders as well as in AD. Tau expression levels depend upon the mouse age, which potentially may also affect its performance in various behavioral tests, although there have been mixed reports on this issue^20^. We studied mice that were 11–13 months old during recordings, which is an age range when substantial tau-related cellular and behavioral pathology have been reported for the htau line^16,20-22^. To the best of our knowledge, the measurements reported in this study are the first to be conducted in the context of exposure to SARS-CoV-2 VLPs.

Our data indicate several interesting findings. First, neuronal activity patterns of htau mice were significantly different than those of WT mice under baseline conditions. The htau mice showed lower spontaneous firing rate levels, in agreement with previous findings using other tau-expressing mouse models^23^; however, their stimulus-evoked activity metrics were higher than those of WT mice (Fig. 2). Second, upon exposure to VLPs, both genotypes show a short-term increase in the measured stimulus-evoked neuronal firing properties, but htau mice also showed changes in spontaneous activity patterns, as well as changes in response to the vehicle injection (Figs. 4-5). This may relate to a differential response of the htau mice to the IP injection and/or the weekly anesthesia and sedation procedure that was part of the imaging sessions. Increased depressive-like behavior and increased measures of anxiety were reported in htau mice^24^, and the stress conditions associated with an IP injection and repeated anesthesia sessions may contribute to this difference between the WT and htau groups. Finally, while WT mice showed long-term increases in the measured stimulus-evoked activity metrics, the htau mice showed a more complex repertoire of altered activity patterns. The measured metrics, both for stimulus-evoked and spontaneous activity, showed both increases and decreases with respect to their baseline levels. We hypothesize that the expression of the wild-type human tau protein in these mice has a major effect that changes the excitability of cortical neurons in the htau mice (Fig. 2D-E). when these mice face a major insult, such as the exposure to VLPs, their functional neuronal circuits are less stable and easier to be altered. Our findings also support the hypothesis that expression of the wild-type tau protein in htau mice makes them more vulnerable to VLP exposure. This may be similar to the increased vulnerability of patients at increased risk of developing or with neurodegenerative conditions to develop symptoms of COVID-19 infection^25,26^ related long COVID (Post-acute Sequelae of SARS-CoV-2 infection).

## Methods

All surgical and experimental procedures were conducted in accordance with protocols approved by the Lerner Research Institute and Oregon Health and Science University Institutional Animal Care and Use Committees and Institutional Biosafety Committees.

### Preparation of VLPs

Plasmid expression vectors encoding each of the SARS-CoV-2 N, M, E, and S proteins were constructed by standard cloning methods, using synthesized codon optimized sequences, as reported^9^. M, E, N, and S plasmids were transfected at a ratio of 5:1:5:1 into suspension-grown Expi293F cells using Expifectamine reagent (ThermoFisher). Culture supernatant was passed through a 0.45 μm filter, followed by ultracentrifugation through a 20% sorbitol cushion at 30,000 rpm for 2h. The pellet was resuspended in 1/100^th^ original volume of phosphate-buffered saline (PBS). VLP structure assessed by transmission electron microscopy of negatively-stained samples.

### Dose-response curve for the effects of VLPs on the Hypothalamic-Pituitary-Adrenal axis and analysis of cytokines and chemokines in plasma and the hippocampus

Nine-month-old C57BL/6J WT mice (*n* = 19) were injected with VLPs (0 [*n* = 5], 0.3 [*n* = 8], or 1 [*n* = 6] μg, IP). Vehicle media without VLPs but otherwise processed the same way was used for the 0 dose injections. One hour later, blood was collected from the mandibular vein in unanesthetized mice with 0.5M EDTA and centrifuged at 40,000 *g* for 10 min at 4°C to collect plasma, as described^27^. Twenty-four hours later, the mice were euthanized by cervical dislocation and plasma and hippocampal cytokines and chemokines levels were analyzed using a Luminex panel (46PLEX, Mouse Magnetic Luminex Assay). Hippocampi were homogenized in a lysis buffer consisting of 1M Tris-Cl, 6M NaCl, 10% SDS, and 0.5M EDTA, 1% Triton-X, and protease inhibitor (Roche, Sigma Aldrich, catalog #11836170001, St. Louis, MO). Total protein amounts were determined with a BCA protein assay kit (Pierce, Thermo Scientific, catalog #23225, Waltham, MA). Plasma corticosterone was analyzed using a radioimmune assay kit (MP Biomedicals, Irvine, CA). Intra-assay coefficient of variation was 10% and the inter-assay coefficient of variation was 7%.

### Surgical preparation of mice

We used C57BL/6J mice and mice that express the human tau protein^16^ (WT and htau mice, JAX strain #000664 and #005491, respectively). All mice were implanted with cranial windows above their left S1 and M1; 10 months old at the time of surgery; *n* = 12 C57BL6/J mice, 6 males and 6 females; n = 6 htau mice, 4 males and 2 females). The window was placed 1 mm anterior to 4 mm posterior to Bregma, 0.5-3.5 mm lateral to midline, similar to previously-published works^28-30^. Mice were first anesthetized using isoflurane (3% for induction, 1.5% during surgery), the scalp skin was cut and removed, and a rectangular craniotomy was drilled (OMNIDRILL35, World Precision Instruments). An adeno-associated virus (AAV) solution expressing jGCaMP7s under the human synapsin promoter (Addgene catalog number 104487-AAV1, diluted 1:10 with saline) was injected into the cortex according to mouse brain atlas coordinates^31^ (Injection locations anterior/lateral to bregma: M1: two locations, 0.5/-1 mm and -0.5/-1.25 mm; S1: two locations, - 0.5/-1.9 mm and -1.75/-2.5 mm, S1 hindlimb and S1 barrel field, S_HL_ and S_BF_ regions, respectively). Forty nL of virus solution were injected per location, 250 and 500 μm under the pia using a 1 mm-diameter pulled and beveled glass micropipette (P-100 puller and BV-10 beveller, Sutter). The craniotomy was covered with a glass window (two-layer cranial window: 5×3×0.17mm^3^ glass glued to a 6×3×0.17mm^3^ glass; Tower Optical) and secured to the skull bone using ethyl cyanoacrylate glue (Krazy Glue) and dental cement (Contemporary Ortho-Jet, Lang Dental). A custom stainless-steel head post was also cemented to the skull using the same dental cement. Mice were given post-operative care for pain management and 3-4 weeks for recovery before the beginning of recordings.

### VLP exposure

A VLP dose of 0.3 μg/mouse was injected intraperitoneally (IP) one hour before the initiation of the recording session on the treatment day. Vehicle mice were injected with an identical volume of the vehicle solution one hour prior to the initiation of the imaging session on the same day.

### Recording of brain activity

Similar to previous works^28,29,32-35^, mice were lightly anesthetized using isoflurane (0.5%), sedated using intramuscular injection of Chlorprothixene Hydrochloride (30 μl of 0.33 mg/ml solution, Spectrum Chemical MFG Corp), and were kept in the dark on a 37°C heating pad for at least 30 minutes before the start of recording. We first recorded spontaneous brain activity using a Bergamo II two-photon microscope (Thorlabs) equipped with a resonant-galvo scanners (1024×1024 pixels, 15 frames/sec, 600×600 μm^2^ field of view, FOV) and a GaAsP PMT detector. jGCaMP7s signal was excited using 950nm light (Insight X3, Spectra-Physics) and a 16x objective (Nikon CFI LWD). The fluorescence signal was collected using a 525/50 emission filter (Chroma). For monitoring spontaneous activity, each FOV was recorded for 200 sec. Typically, two FOVs were recorded from each of the S1 and M1 regions of each mouse. Then, a thin needle electrode was attached to mouse hindlimb (29-gauge needle electrode, AD Instruments) and connected to a pulse stimulator (A-M Systems, model 2100). The recording was divided into 10 cycles. Each cycle started by 5 sec of baseline recording, followed by 5 sec of paw stimulation with 25 electric stimuli. Each stimulus consisted of a 1.5mA square amplitude pulse of 100msec duration, which was followed by 100msec without stimulus, for a total of 5 sec. The recording cycle was concluded by acquiring additional 5 sec of brain activity without stimulation. A delay of 5 sec without recording separated subsequent cycles. Typically, 2 FOVs per brain region were recorded from one M_1_ spot and the S_HL_ region from each mouse in all recording sessions. Mice were recorded for 2-3 baseline recording sessions before they were exposed to VLP/vehicle. Then, they were recorded on the the exposure day, with the recording session starting 1 hour after the IP injection. Afterwards, the mice were recorded once a week for additional 6 weeks. All recording sessions were identical.

### Brain activity data analysis

Analysis was based upon custom MATLAB scripts (MathWorks), similar to our previously-published data^28,34,35^. Raw fluorescence movies were registered using the TurboReg plug-in in ImageJ to correct small movements^33,36^. Regions of interest (ROIs) corresponding to all identifiable somata were labeled using CellPose software^37^. Recorded data were inspected for low expression of jGCaMP7s, movement artifacts, or low image quality, and we excluded recorded movies with such issues from the analysis. We note that the htau mice express low level of background GFP signal, and FOVs where the contrast of the somatic jGCaMP7s signal with respect to the surrounding GFP signal were also excluded from the analysis. Fluorescence signal from all pixels of each ROI were averaged to calculate the time-dependent fluorescence signal (F) and were corrected for neuropil contamination^38^ using r=0.7, baseline signal (F_0_) for each neuron was estimated as the mean value of F three seconds prior to the initiation of the paw stimulation for stimulated data, or as the median fluorescence value for spontaneous recordings. The single-cell fluorescence change was calculated using the formula: ΔF/F_0_= (F-F_0_)/F_0_. Action potential (AP) firing number and timing were estimated for spontaneous activity using the SpikeML algorithm^39^ that extracted these values from the single cell ΔF/F_0_ using the following jGCaMP7s parameters^32^ (1AP amplitude=35%, decay time=600 msec, Hill coefficient=2.49). SpikeML output was used to calculate the fraction of active neurons (neurons with at least one identified AP during recording) and the average cellular firing rates and to compare changes across recording sessions. For analyzing short-term changes, we compared measurements from baseline (mean of recordings for mice with more than one baseline recording), the exposure week, and the following week, while for identifying long-term changes, we averaged the measured values across the early time after exposure (same day and the following week), middle (weeks 2-3 after exposure), and late (weeks 4-6 after exposure).

For quantifying the stimulated neuronal activity, we calculated the following values: F_base_ – average fluorescence during the 3sec before each paw stimulation started. F_stim_ - average fluorescence during the 5sec of paw stimulation plus the following 1 second to compensate for jGCaMP7s slow kinetics.

F_decay_ - average fluorescence during the 3 sec starting 1 sec after the end of paw stimulation until 4 sec after the end of the paw stimulation. For recording of each FOV, we ran 10 cycles of paw stimulation, with 5 sec of 5 stimuli/sec in each, therefore, F_base_, F_stim_, and F_decay_ were vectors containing 10 values each. For defining whether a neuron is tuned, we ran a paired Student’s t-test (*p* = 0.01) between the F_base_ and F_stim_ vectors. For defining whether a neuron has sustained activity, we ran a paired Student’s t-test (*p* = 0.01) between F_base_ and F_decay_. A cell that passed as significant in both tests was defined as a neuron with both tuned and sustained activity. Next, we defined the median area under the curve (AUC) as the median of the sum of the ΔF/F_0_ values during 6 sec in each of the 10 stimulus cycles (5 sec of each stimulus cycle + additional 1 sec after the end of each cycle). This value was divided by 2 and then we subtracted the median of the sum of ΔF/F_0_ values during the 3 sec of baseline period prior to the first stimulus.

## Statistical analysis

Statistical analysis was conducted using GraphPad Prism (version 10.1.2). For the Luminex and plasma corticosterone analysis, one-way ANOVAs with Dunnett’s *post hoc* tests were used. To assess relationships between hippocampal chemokine levels, Pearson’s correlations were used. For analyzing the repeated neuronal activity measurements, mixed-effect models with restricted maximum likelihood and Tukey Honest Significant Difference *post hoc* tests (p≤0.05) were used to identify changes between groups (VLP and vehicle) and time points. Data from WT and htau mice were analyzed separately due to the changes among these groups that were apparent during baseline recordings. Significant *post hoc* comparisons for WT mice were shown only for comparing the same brain region across different times or for comparing the VLP and vehicle groups for the same brain region. For htau mice, we did not perform *post hoc* comparisons due to the low number of tested mice per group.

## Acknowledgements

The authors would like to thank Dr. Christopher Nelson for helpful comments and suggestions to improve the manuscript. The authors would like to thank the HHMI Janelia GENIE team for sharing the jGCaMP7s indicator, Dr. Vanzetta and their team for sharing the SpikeML algorithm, and Dr. Pachitariu and their team for sharing the CellPose software. JR and HD were supported by grants R21AG065914 and R21AG065914-01S1from the NIH/NIA.

## Notes

### Competing Interest Statement

The authors have declared no competing interest.

